# Patterns of ‘Analytical Irreproducibility’ in Multimodal Diseases

**DOI:** 10.1101/2020.03.22.002469

**Authors:** Abigail R Basson, Fabio Cominelli, Alex Rodriguez-Palacios

## Abstract

Multimodal diseases are those in which affected individuals can be divided into subtypes (or ‘data modes’); for instance, ‘mild’ vs. ‘severe’, based on (unknown) modifiers of disease severity. Studies have shown that despite the inclusion of a large number of subjects, the causal role of the microbiome in human diseases remains uncertain. The role of the microbiome in multimodal diseases has been studied in animals; however, findings are often deemed irreproducible, or unreasonably biased, with pathogenic roles in 95% of reports. As a solution to repeatability, investigators have been told to seek funds to increase the number of human-microbiome donors (N) to increase the reproducibility of animal studies (doi:10.1016/j.cell.2019.12.025). Herein, through simulations, we illustrate that increasing N will not uniformly/universally enable the identification of consistent statistical differences (patterns of analytical irreproducibility), due to random sampling from a population with ample variability in disease and the presence of ‘disease data subtypes’ (or modes). We also found that studies do not use cluster statistics when needed (97.4%, 37/38, 95%CI=86.5,99.5), and that scientists who increased N, concurrently reduced the number of mice/donor (*y*=-0.21x, *R^2^*=*0.24*; and vice versa), indicating that statistically, scientists replace the disease variance in mice by the variance of human disease. Instead of assuming that increasing N will solve reproducibility and identify clinically-predictive findings on causality, we propose the visualization of data distribution using kernel-density-violin plots (rarely used in rodent studies; 0%, 0/38, 95%CI=6.9e-18,9.1) to identify ‘disease data subtypes’ to self-correct, guide and promote the personalized investigation of disease subtype mechanisms.

**Highlights:**

- Multimodal diseases are those in which affected individuals can be divided into subtypes (or ‘data modes’); for instance, ‘mild’ vs. ‘severe’, based on (unknown) modifiers of disease severity.
- The role of the microbiome in multimodal diseases has been studied in animals; however, findings are often deemed irreproducible, or unreasonably biased, with pathogenic roles in 95% of reports.
- As a solution to repeatably, investigators have been told to seek funds to increase the number of human-microbiome donors (N) to increase the reproducibility of animal studies.
- Herein, we illustrate that although increasing N could help identify statistical effects (patterns of analytical irreproducibility), clinically-relevant information will not always be identified.
- Depending on which diseases need to be compared, ‘random sampling’ alone leads to reproducible ‘patterns of analytical irreproducibility’ in multimodal disease simulations.
- Instead of solely increasing N, we illustrate how disease multimodality could be understood, visualized and used to guide the study of diseases by selecting and focusing on ‘disease modes’.

## Introduction

Multimodal diseases are those in which affected individuals can be divided into subtypes (or ‘data modes’); for instance, ‘mild’ vs. ‘severe’, based on (unknown) modifiers of disease severity. Since the availability of DNA-sequencing platforms, there have been major advances in our understanding of the human microbiome, its ecological complexity, and temporal oscillations. However, to differentiate the causal connection between microbiome alterations and human diseases (from that of secondary alterations due to disease), animal models, primarily germ-free rodents transplanted with human gut/fecal microbiota (**hGM-FMT**), have been critical as *in vivo* phenotyping tools for human diseases. Unfortunately, despite considerable efforts from organizations and guidelines to help scientists design and report preclinical experiments (*e.g*., ARRIVE)^1,2^, there are still concerns of study reproducibility.

Studies have described novel technical sources of ‘artificial’ microbiome heterogeneity that could explain why hGM-FMT study results vary^2–6^. In our own work^2^, we discovered that scientists lacked appropriate methods for the description and analysis of cage-clustered data. To help scientists to self-correct issues on rodent experimentation, we identified ‘six action themes’ and provided examples, and statistical code, on how to use and compute ‘study power’ as a reproducible parameter that could enable inter-laboratory comparisons and improve the planning of human clinical trials based on preclinical data^2^.

In this regard, a recent perspective article on hGM-associated rodent studies by Walter *et al*.^7^ (“*Establishing or Exaggerating Causality for the Gut Microbiome: Lessons from Human Microbiota-Associated Rodents*”; published in Cell, January 23^rd^ 2020) recommended to scientists seek additional funding to increase the number of human donors (**N**) as a main solution to improve experimental rigor and reproducibility, and to determine the causal role of the hGM in disease. Given that large disease variability is experimentally problematic for both humans and animals, we hypothesized that increasing N would not ensure consistent results due to the aleatory effects of random sampling of subjects from a population with multimodal disease distributions (*i.e*., multimodal: >2 types of modes or ‘subtypes of disease data’ can be seen in a population; the most, and the least diseased). To verify this hypothesis in the context of hGM and N, we used published (observed) preclinical distribution (disease variability) estimates to conduct a statistical and visualization analysis of the impact of repeated random sampling on the significance of statistical comparisons between simulated disease groups, at various N.

Underscoring the importance of the central limit theorem (which can be visualized in[8]), simulations indicate that more studies addressing disease multimodality (independent of N; personalized disease subtyping studies) are preferable than fewer studies with larger N that do not address disease multimodality. After examining the statistical content of 38 studies^8–45^ listed in Walter *et al*,^7^ we found that scientists who increased N, concurrently reduced the number of MxD, indicating that statistically, scientists replace the disease variance in mice by the disease variance in humans in their hGM-FMT studies. Further, studies lacked proper clustered-data statistics to control for animal density; which is a major source of misleading results (false-positive, or false-negative), especially when scientists prefer to house many rodents per cage, and when the number of mice per experiment is low^2,46^.

Herein, we provide a conceptual framework that illustrates various patterns of analytical irreproducibility by simulating and integrating the dynamics of: N, random sampling, group means, sample variance, and the population disease diversity that could be visualized as unimodal, bimodal or multimodal, through the use of kernel-based violin density plots for the identification of data subtypes. Simulations and provided examples could help scientists ***i)*** visualize the dynamics of random sampling from a heterogeneous population of healthy and diseased subjects, ***ii)*** decide on N once preclinical data are generated, and ***iii)*** improve experimental rigor in hGM-FMT studies.

## RESULTS

### ‘Disease subtypes’ occur in simulations using published data and UNIMODAL distributions

In microbiome rodent studies, the selection of a sufficient number of both human donors (N), and the number of mice required to test each human donor (**MxD**), is critical to account for the effects of random sampling, which exist when the hGM induces variable disease severity in humans and rodents. Thus, to visualize the variability of disease severity (data subtypes/modes) in rodents, and the effect of N on the reproducibility of pairwise statistical comparisons between groups of hypothetically, randomly selected human donors, we first conducted a series of simulations using the mean±SD (observed data) from hGM-FMT mice in Basson *et al*.^46^ (note the dispersed overlapping variances, SD in **Figure 1A**). Using the observed data we generated random datasets using functions designed to draw numbers from an inversed Gaussian distribution (with unimodal normal continuous probability; 0,∞). We demonstrate how the random selection of donors (sampled as groups for each of three iterative datasets) influence the direction and significance level in pairwise comparative statistics (**Figure 1B**).

**Figure 1.**
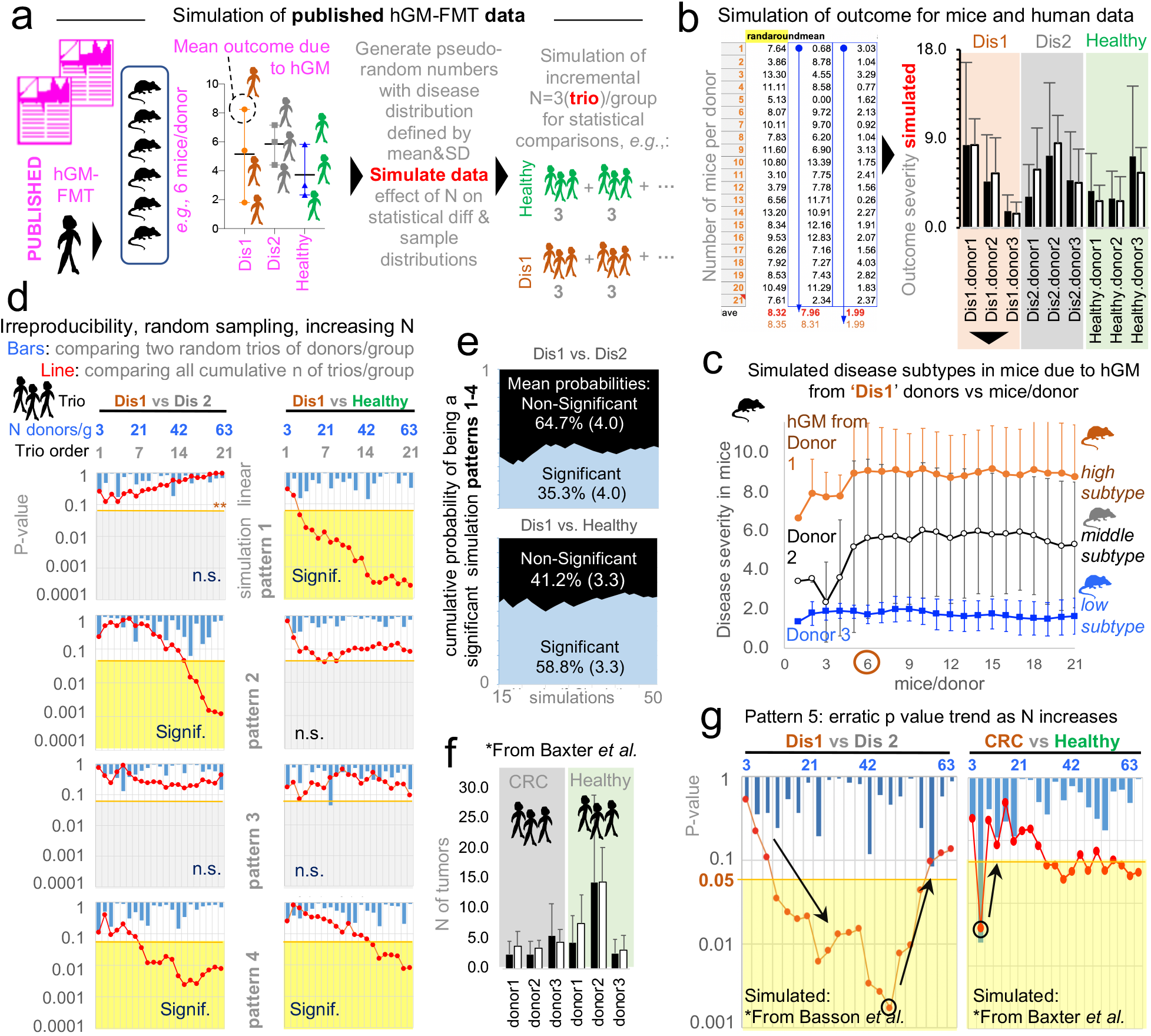
Random sampling from overlapping diseases yield ‘linear patterns of analytical irreproducibility’. Simulations on observed data from Basson *et al*^46^ to visualize naturally/highly variable disease/healthy datasets. **a**) Method overview to generate pseudo-random numbers and simulations from published (observed) data. **b**) Visualization of simulated outcome from random integers generated based on 3 donors/group for Disease 1 (‘Dis1’), Disease 2 (‘Dis2’), and healthy groups. **c**) Simulation of hGM transplanted into mice yields reproducible simulated ‘disease data subtypes’ from 6 mice/group. **d**) Four patterns of analytical irreproducibility. Representative simulations comparing 2 groups of donors, with N ranging from 3 (trio)donors/group to 63, in multiples of 3 (cumulative addition of new trios per group). Y axis, p-value of differences using 2-group Student-t test. Notice as N increases, the cumulative significance (red line) exhibit different linear patterns due to variance introduced by random sampling. **e**) Cumulative probability of a simulation to yield a significant difference (blue; significant, black; non-significant; parentheses, std. dev.). A comparison was deemed significant, if at least one p-value<0.05 across simulations with N between 3 and 63 donors/group. **f**) Visualization of simulated outcome using observed data from Baxter *et al*^16^. **g**) Random simulations illustrate two other possible analytical patterns. Notice as N increases, group differences become more significant, until an inflection point, where adding more donors makes the significance disappear. See **Supplementary Figure 1** for additional examples and computed R^2^ value to illustrate the linearity of the correlation between N and statistical significance.

Simulations showed that the number of MxD is important because mice have various response patterns to the hGM (*i.e.,* disease severity, data subtypes/modes), which can be consistently detected depending on the MxD and thus the variability introduced by random sampling. Simulations showed that for the three group datasets (plotted as ‘Dis1’, ‘Dis2’ and ‘Healthy’), it was possible to reproducibly identify two-to-three unique donor disease severity subtypes (data modes) in mice induced by the hGM (‘high’, ‘middle’, and ‘low’ disease severity). Simulation plots made it visually evident that testing <4-5 MxD yielded mean values more likely to be affected by intrinsic variability of random sampling; thus, making studies with >6 MxD more stable and preferable. Conversely, studies with 1-2 MxD are at risk of being strongly dependent on randomness. Iterative simulations showed that the mean effect (*e.g*., ileal histology) in transplanted mice varies minimally (*i.e*., stabilizes) after 7±2 MxD, depending on the random dataset iterated. Beyond that, increasing MxD becomes less cost-effective/unnecessary if the focus is the human donors (**Figure 1C**).

### Random sampling from overlapping diseases yield ‘linear patterns of analytical irreproducibility’

Often, published literature contains figures and statistical analysis conducted with 3 donors per disease group. Thus, to mimic this scenario and to examine the role of random sampling on the reproducibility of pairwise statistical results (‘significant’ vs. ‘non-significant’), we conducted, ***i)*** multiple 3-donor/group (‘trio-trio’) pairwise comparisons, and ***ii)*** a simultaneous overall analysis for the cumulative sum of all the 3-donor trios simulated for each disease group. That is, we monitored and quantified whether results for each random iteration were significant (using univariate Student’s t-statistics p<0.05) or non-significant (p>0.05) for groups of simulated donor datasets (‘Dis1’ ‘Dis2, and ‘Healthy’). Assessing the effect of random sampling at various N, and also as N accumulated, we were able to illustrate that pairwise trio-trio comparisons between the simulated datasets almost always produce non-significant results when iterative trios were compared (due to large SD overlapping; see bars in **Figure 1D** representing 21 sets of pairwise trio-trio p-values). However, as N increases by the cumulative addition of all (mostly ‘non-significant’) donor trios (*i.e*., N increases in multiples of 3, for a range of N between 3 and 63 donors/group; [3, 6, 9, 12…63]), pairwise statistical comparisons between the simulated datasets did not produce consistent results (see line plots in Figure 1D representing p-value for cumulative addition of donors when sampling iterations were simulated).

Results are clinically relevant because the simulated N, being much larger (63 donors/group) than the largest N tested by one of the studies reviewed by Walter *et al*^7^ (21 donors/group)^40^ demonstrates that the analysis of randomly selected patients would not always yield reproducible results due to the chance of sampling aleatory sets of individuals with varying degrees of disease severity, regardless of how many donors are recruited in an study. To provide a specific example, using ‘Dis1’ as a referent, cumulative pairwise comparisons (vs. ‘Dis2’, and vs. ‘Healthy’) revealed at least five different patterns of ‘irreproducible’ statistical results as N increased between 3 and 63 per group. **Figure 1D** illustrates four of these variable cumulative linear patterns of analytical irreproducibility, in which, remarkably, ***i)*** ‘Dis1’ becomes significantly different vs. Dis2, and vs. ‘Healthy’, as N increases, ***ii)*** ‘Dis1’ becomes significantly different from ‘Dis2’ but not vs. ‘Healthy’, ***iii)*** ‘Dis1’ was significantly different from healthy but not vs. ‘Dis2’, and ***iv)*** ‘Dis1’ never becomes significantly different despite sampling up to 63 donors/group. See **Supplementary Figure 1** for complementary plots illustrating linearity of patterns (R^2^, mean 0.51±0.23, 21 simulations) Hence, results clearly illustrate that seeking funds to recruit more donors is not a prudent statistical solution to the problem of understanding disease causality of widely variable conditions in both humans and animals. By analytical irreproducibility, herein, we refer to the inability to reproduce the direction and statistical significance of a test effect when analyses are conducted between groups created by the random selection of subjects from distributions defined based on observed (mean±SD) data.

### 100,000 Monte Carlo simulations illustrate the effect of randomness on analytical reproducibility

To summarize the overall significance of the inconsistent patterns observed via random sampling, we computed an aggregate ‘cumulative probability of being a significant simulation’ for 50 pairwise statistical simulation sets fulfilling the 4 linear patterns described above. Emphasizing the concept that increasing N is not a reproducible solution, **Figure 1E** shows that only 35.3±4.0% of comparisons between ‘Dis1’ and ‘Dis2’, and 58.8±3.3% for ‘Dis1’ and ‘Healthy’ were significant.

Expanding the validity of these inverted-Gaussian simulations for N=63 donors/group, we then conducted ***i)*** Monte Carlo adjusted Student’s unpaired t-tests, and ***ii)*** Monte-Carlo adjusted one-way ANOVA with Tukey correction for family errors and multiple comparisons. Monte Carlo simulations used data drawn from a normal (non-inverted) Gaussian distribution around the group means with a pooled SD of ±4, and were conducted using GraphPad, a popular statistical software in published studies. To estimate a probability closer to the real expectation (narrower confidence intervals), 100,000 simulations were performed. Supporting the observations above (based on inverse normal simulations), Monte Carlo Gaussian simulations showed that, using pairwise comparison, ‘Dis1’ would be significantly different from Dis 2 (adjusted T-test p<0.05) only 57.7% of the time (95%CI=58-57.4), with 1540 simulations producing negative (contradictory) mean differences between the groups. Compared to ‘Healthy’, ‘Dis1’ and ‘Dis2’ were significant only 9.1% (95%CI=9.2-8.9) and 78.3% (95%CI=78.6-78.1) of the time, respectively.

Under the ‘Weak Law of Large Numbers’^47–49^, and randomization principles, it is almost always possible to detect some level of statistical significance(s) and mean group differences when asymptotic mathematical methods based on numerous simulations are used, for example, as a surrogate for multiple experiments which are not possible in real research settings. However, in this case, the mean simulated differences yielding from 100,000 simulations were minuscule (1.6 for ‘Dis1’-‘Dis2’; −1.97 ‘Healthy’-‘Dis2’, and 0.42, ‘Healthy’-‘Dis1’). Compared to the range of disease variance for each disease, such minuscule differences may not be clinically relevant to explain disease variance at the individual level. Note that the SD was 4, therefore it is intuitive to visualize in a numerical context such as small differences across greatly overlapping unimodal simulations. Correcting for family errors, One-way ANOVA corrected with 10,000 Monte Carlo simulations with N=63/group, showed that at least one of the three groups would be statistically different in approximately only 67.2% of the simulations (95%CI=64.2-70.0), whereas in 32.8% (95%CI=64.2-70.0) of simulations, the groups would appear as statistically similar (see **Table 1** for estimations after 100,000 Monte Carlo simulations; note narrower CI as simulations increase). The comparison of ‘Dis1’ vs. ‘Dis2’ in Table 1, clearly demonstrates that the percentage of cases, in which a simulation could be significant, depends on the degree of data dispersion. For example, simulations with SD of 4, compared to SD of 10, produce significant results less often, illustrating how data with larger dispersions contribute to poor statistical reproducibility, which cannot necessarily be corrected by increasing N.

**Table 1.** Comparative percentages of simulations that yielded significant results for two statistical approaches based on randomly simulated data derived from unimodal distributions

|  | Inverse Normal Gaussian | Monte Carlo Normal Gaussian |  |
| --- | --- | --- | --- |
| Simulation, n= and statistical test | 50 T-tests (significant cumulative linear pattern*) (95%CI=) | 100,000 Adjusted T-tests (overall significance with N=63/group)(95%CI=)<sup>a</sup> | 100,000 Adjusted One Way with multiple comparison Tukey test<sup>b</sup> |
| Dis1 vs Dis2 | 35.3% (22.9, 50.8) | 57.7% (57.4, 58.0) | 37.8% (37.5, 38.1) |
| Dis1 vs Healthy | 58.8% (43.2, 71.8) | 9.1% (8.9, 9.3) | 3.8% (3.7, 3.9) |
| Dis2 vs Healthy | ND | 78.3% (78.0, 78.6) | 59.6% (59.3, 59.9) |
|  |  |  | One Way ANOVA<br>p<0.05 68.1% (68.4 to 67.8)<br>p>0.05 31.9% (32.2 to 31.6) |

\*Not overall p-value at N=63. ND, not determined.
<sup>a,b</sup> Notice that the percentage of simulations achieving significance is inflated when analysis for three groups is conducted with T-tests (instead of ANOVA) which does not control for false positives due to family errors. Proper comparison between >2 groups should be performed with methods to control for such family errors (e.g., ANOVA-post-hoc Tukey statistics). Note that the percentage is different as illustrated in Figure 1D because the patterns with non-linear behavior are not considered.

### Random sampling can lead to ‘erratic patterns of analytical irreproducibility’ as N increases

To increase the external validity of our observations, we next simulated the mean±SD data published from a hGM-FMT study on colorectal cancer conducted by Baxter *et al*^16^. In agreement with Basson *et al*, Baxter *et al* revealed comparably bimodal colorectal cancer phenotypes in mice resulting from both the diseased (colorectal cancer) patients and healthy human donors (**Figure 1F**). Equally important, we observed for both Basson *et al*^46^ and Baxter *et al*^16^, what we describe as the fifth ‘pattern of analytical irreproducibility’ in this report. That is, in some cases, the steady addition of donor trios/group (as simulations proceeded for increasing values of N) made it possible to identify simulations where erratic changes in the statistical significance for group comparisons switched randomly, yet gradually, from being significant to non-significant as more donor trios were ‘recruited’ into the simulations (**Figure 1G**). Clinically relevant, simulations indicated (in a reproducible manner), that adding extra patients could at times actually invert the overall cumulative effect of the p-value, possibly due to the variable distribution and multimodal nature of the human and rodent responses to experimental interventions. As such, simulations indicate that it is advisable to conduct several *a-priori* determined interim results in clinical trials to ensure that significance is numerically stable (p<0.05), as well as the relevance of personalized analysis to examine disease variance in populations. Unfortunately, there are no guidelines or examples available to assist in determining how many donors would be sufficient, and to visualize the effect of random sampling of individuals from a vastly heterogeneous population of healthy and diseased subjects, once rodent preclinical data is generated.

### Violin plots and statistical methods for visualization of MULTIMODAL ‘disease data subtypes’

To visualize the underlying mechanisms that could explain the ‘linear and erratic patterns of analytical irreproducibility’ introduced by random sampling, we first used dot plots based on observed and simulated data, followed by kernel-based statistics and plots. Plot appearance and one-way ANOVA statistics showed that when N is increased, significant results, when present for largely overlapping phenotypes, are primarily due to small differences between sample means (**Figure 2A-B**). Simulations that compared 3 groups of 65 donors/group almost always yielded a significantly different group; however, dot plots show that the significant differences between means are just a small fraction of the total disease variability as verified with Monte Carlo simulations above. That is, as N increases, comparisons can become significant (see plot with 65 donors in **Figure 2C**). In this context, a significant difference of such a narrow magnitude may not be clinically relevant, or generalizable, to explain the presence of a disease phenotype in a population, especially for those individuals at the extreme ranges of the disease distribution.

**Figure 2.**
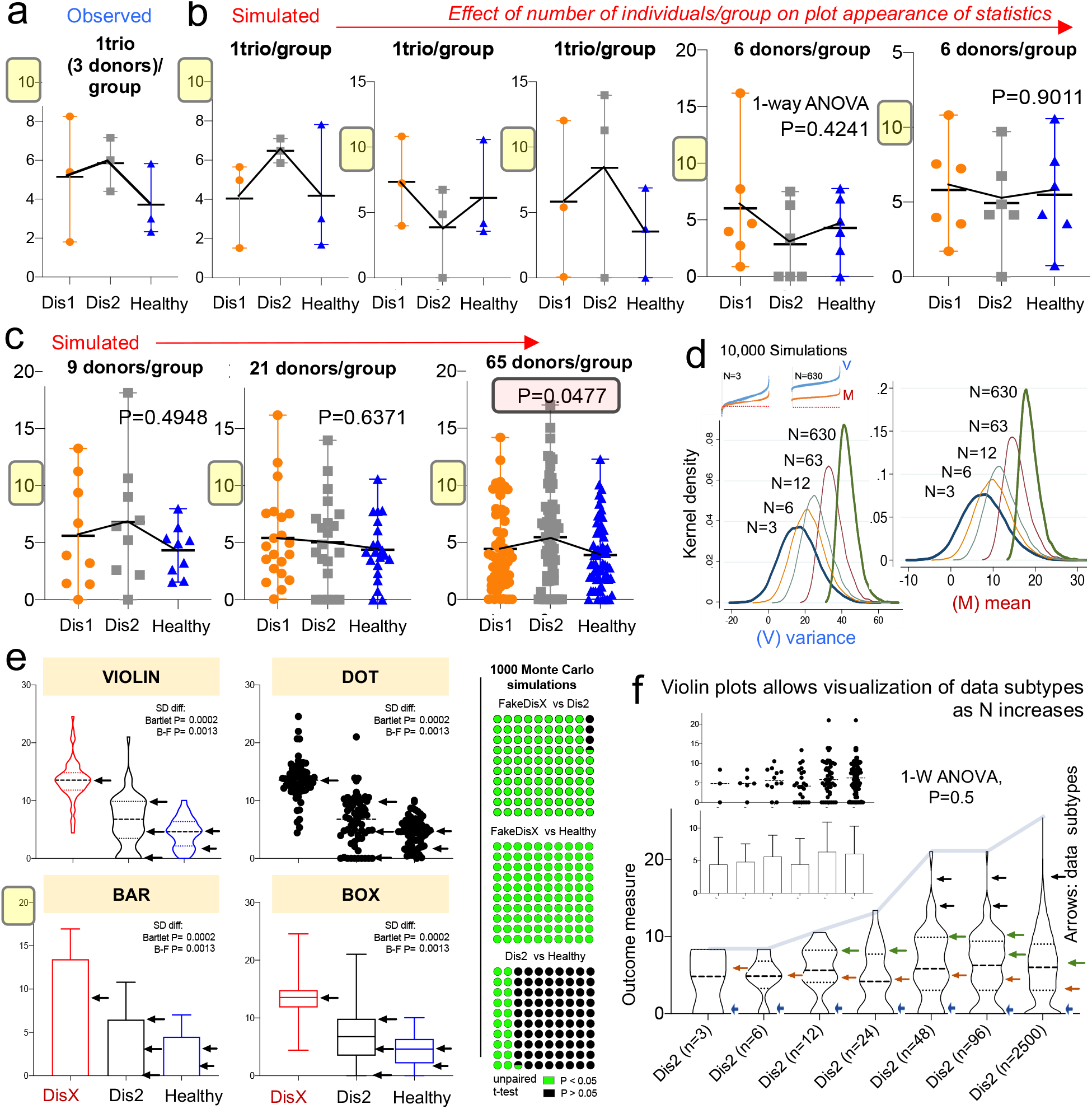
Violin plots enable visualization of data subtypes in simulations of random sampling as a function of N. Observed raw data derived from Basson *et al* published data. **a-b)** Dot plots (mean, range) of observed (1 trio; 3 donors/group), and simulated data (3 and 6 donors/group; panel B). Note that differences are not significant because of the variability between diseases. **c)** Dot plots (mean, range) of simulated data for 9, 21 and 65 donors per group. Note that simulated mean effects became significant with 65 donors/group. However, the mean difference is small compared to the variance of the groups and the difference is not biologically different because it is a function of the total variance (23%). **d)** Kernel density simulations (10,000) based on observed (n=3) and simulated data. Note that as N increases the mean becomes more narrow while the variance widens. See 100,000 Monte Carlo simulations in **Table 1**. **e)** Comparison of visual appearance and data display for simulated data to illustrate ‘disease data subtypes’ in the population (i.e., ‘disease data shoulders’/subtypes shown as arrows). **f)** Violin plots allow visualization of data subtypes as N increases (‘disease data shoulders’/subtypes shown as arrows).

Mechanistically, the detection of significant comparisons can be attributed to the effect that ‘increasing N’ has on the data mean and variance, which increases at a higher rate for the variance as shown in **Figure 2D**. Instead of increasing N as a general solution, we propose to scientists to use violin plots, over other plots commonly encouraged by publishers^50^ (*e.g*., bar, boxplot and dot plots), because violin plots provide an informative approach, at the group-sample level, for making inferences about ‘disease data subtypes’ in the population (see ‘subtypes’ shown with arrows in **Figure 2E**).

Violin plots are similar to a box plot, as they show a marker for the data median, interquartile ranges, and the individual data points^51^. However, as a unique feature, violin plots show the probability density of the data at different values, usually smoothed by a kernel density estimator. The idea of a kernel average smoother is that within a range of aligned data points, for each data point to be smoothened (X0), there is a constant distance size (λ) of choice for the kernel window (radius, or width), around which a weighted average for nearby data points are estimated. Weighted analysis gives data points that are closer to X0 higher weights within the kernel window, thus identifying areas with higher data densities (which correspond to the disease data modes). As an example of the benefits of using violin plots, **Figure 2F** illustrates that as N increases, so does the ability of scientists to subjectively infer the presence of disease subtypes. To strengthen the reproducibility of ‘subtype’ mode identification, herein we recommend the use of statistical methods to identify disease data modes (*e.g*., see the *modes* function in **Methods** and **Discussion**), because as N increases, the visual detection of modes becomes increasingly more subjective as shown in **Figure 2F**.

### Kernel density violin plots help guide subtype analysis to identify biologically significant results

Violin plots and kernel density distribution curves in **Figure 3** illustrate why comparing groups of randomly sampled individuals may not yield biologically relevant information, even though statistical analysis identifies that the mean values differ between compared groups. **Figure 3A** illustrates the different patterns of potential donor subtypes (*i.e*., data modes, visualized in violin plots as disease data/curve ‘shoulders’) that would yield significant results in a single experiment depending on the donors sampled. However, the kernel density plots in **Figure 3B** show that significant findings do not necessarily indicate/yield clinically relevant thresholds or parameters to differentiate between the populations (due to the overlapping and inflation of data ‘shoulders’ in some subjects within the samples). To contrast the data simulated from Basson *et al*., we replaced data from ‘Dis1’ dataset with a Gaussian distributed sample of random numbers (within 13.5±3.5, labeled as ‘fake disease X’; vs. 6.4±4.3, and 4.5±2.5 for ‘Dis2’ and ‘Healthy’, respectively) to illustrate how a kernel plot would appear when significant differences have a clinically relevant impact in differentiating disease subtypes (**Figure 3C-D**).

**Figure 3.**
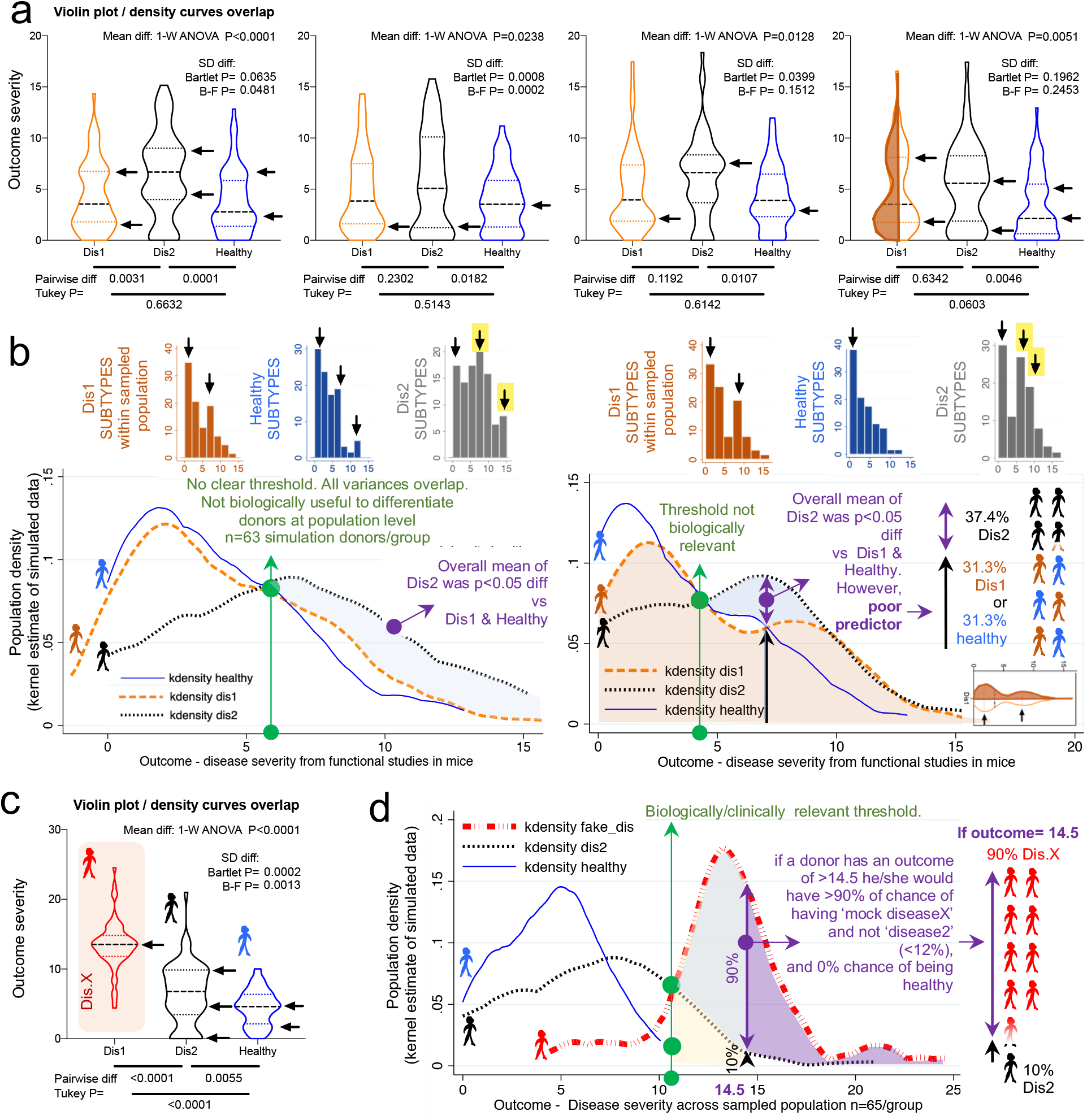
Violin plots illustrate that statistical differences with large N may not have clinical predictive utility at individual level. Violin and kernel plots illustrate statistical vs. biologically relevant differences. **a)** Violin plots of four simulated random number sets illustrate that each set of donors may have unique subtypes of disease illustrated with arrowheads (disease severity scores with higher number of simulated donors). Arrows indicate ‘disease data shoulders/subtypes’ vary with every simulation of 63 donors/group. **b)** Kernel density curves illustrate large overlap of sample population from simulated data (see panel A). Significant differences are highlighted by shaded area. Note the threshold does not have distinctive separation for the plots indicating that it is not biologically useful as a predictor of outcome. **c)** Violin plots illustrate meaningful statistical difference for population (compared to panel 3b). ‘Fake disease X’ (‘DisX’) was generated as a ‘mock’ disease following Gaussian distribution around the mean. Monte Carlo simulations were significant 96.5% (upper limit 97.6, lower limit 95.4%). **d**) Kernel density curves of panel 3c illustrates example of distribution separation with both statistical difference and biological relevance.

Collectively, simulations indicate that the uneven random sampling of subtypes across a disease group would be an important factor in determining the direction of significance if studies were repeated, owing primarily to the probability of sampling data ‘shoulders’ or ‘valleys’ in both healthy and diseased populations.

### Simulation of BIMODAL diseases illustrate mechanism of analytical irreproducibility

In our report thus far, we have used unimodal simulations to show how random sampling affects statistical results. However, there has been an increased interest in understanding data multimodality in various biological processes^52,53^ for which new statistical approaches have been proposed. Methods to simulate multimodal distributions are however not trivial, in part due to the unknown nature of multimodality in biological processes. To facilitate the understanding of the conceptual mechanisms that influence the effect of data multimodality and random sampling on statistical significance, **Figure 4** schematically contextualizes the statistical and data distribution principles that can interfere with reproducibility of statistical results when simulations are repeated.

**Figure 4.**
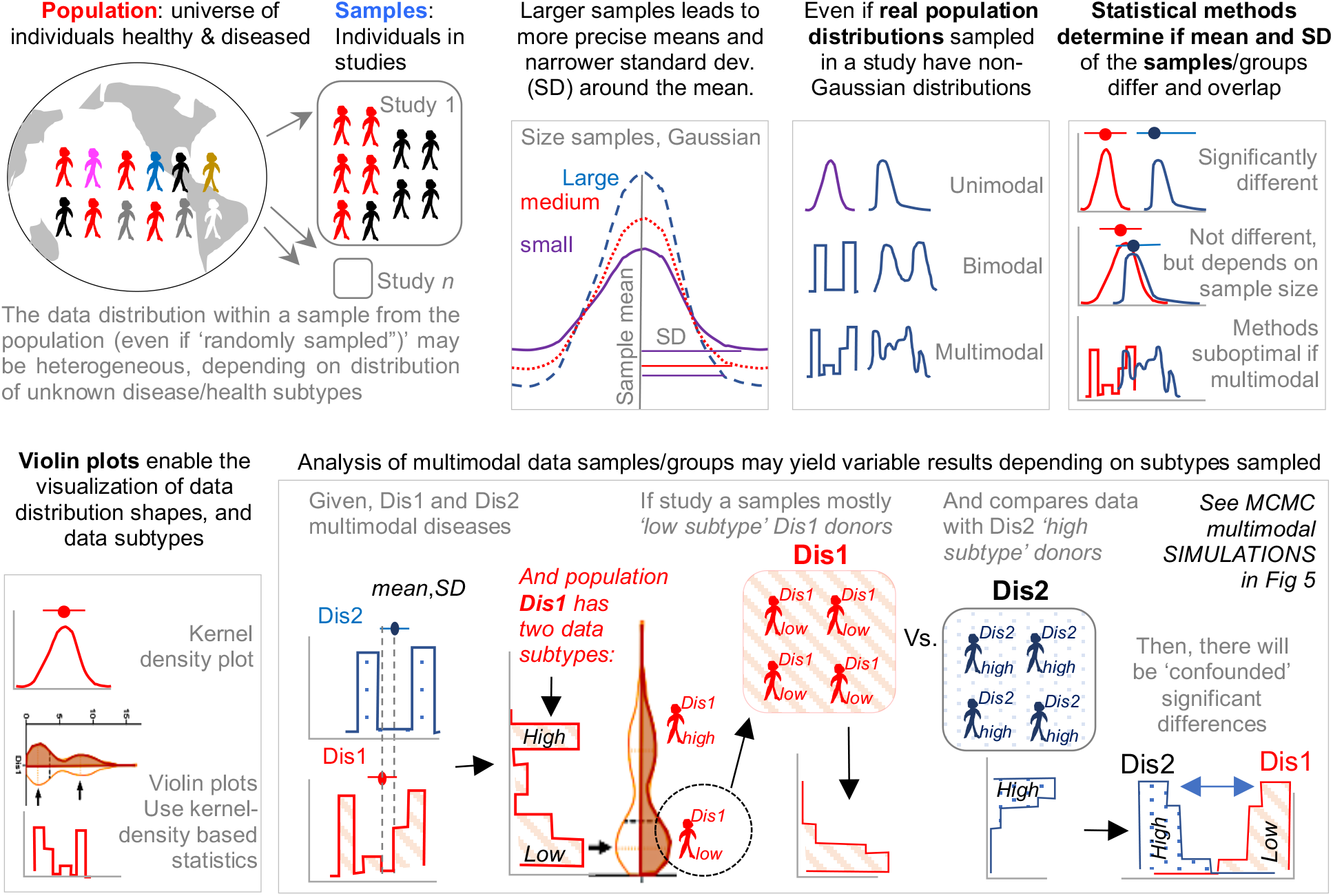
Conceptual framework of the effect of random sampling from a multimodal disease population on the reproducibility of study results. Schematic conceptualization of random sampling from a bi/multi-modal disease distribution (subtypes) and utility of violin and kernel density plots to visualize disease subtypes.

Random simulations from unimodal distributions work on the assumption that numbers (*e.g*., donors’ disease severity) are drawn from a population, *independently* from one another. That is, the probability of sampling or drawing a number from a population is not influenced by the number that was selected prior. While this form of random sampling is very useful in deterministic mathematics, it does not capture the *dependence* of events that occur in biology. That is, in biology, the probability of an event to occur *depends* on the nature of the preceding events. To increase the external validity of our report, we thus conducted simulations based on three strategies to draw density curves resembling bimodal distributions. **Figure 5A** depicts distributions derived from both ‘truncated beta’, and the combination of two ‘mixed unimodal’ distribution functions (*e.g*., two *independent* Gaussian curves in one plot), which are illustrative of multimodality, but not necessarily reliable methods to examine the effects from *dependent* random sampling in multimodality.

**Figure 5.**
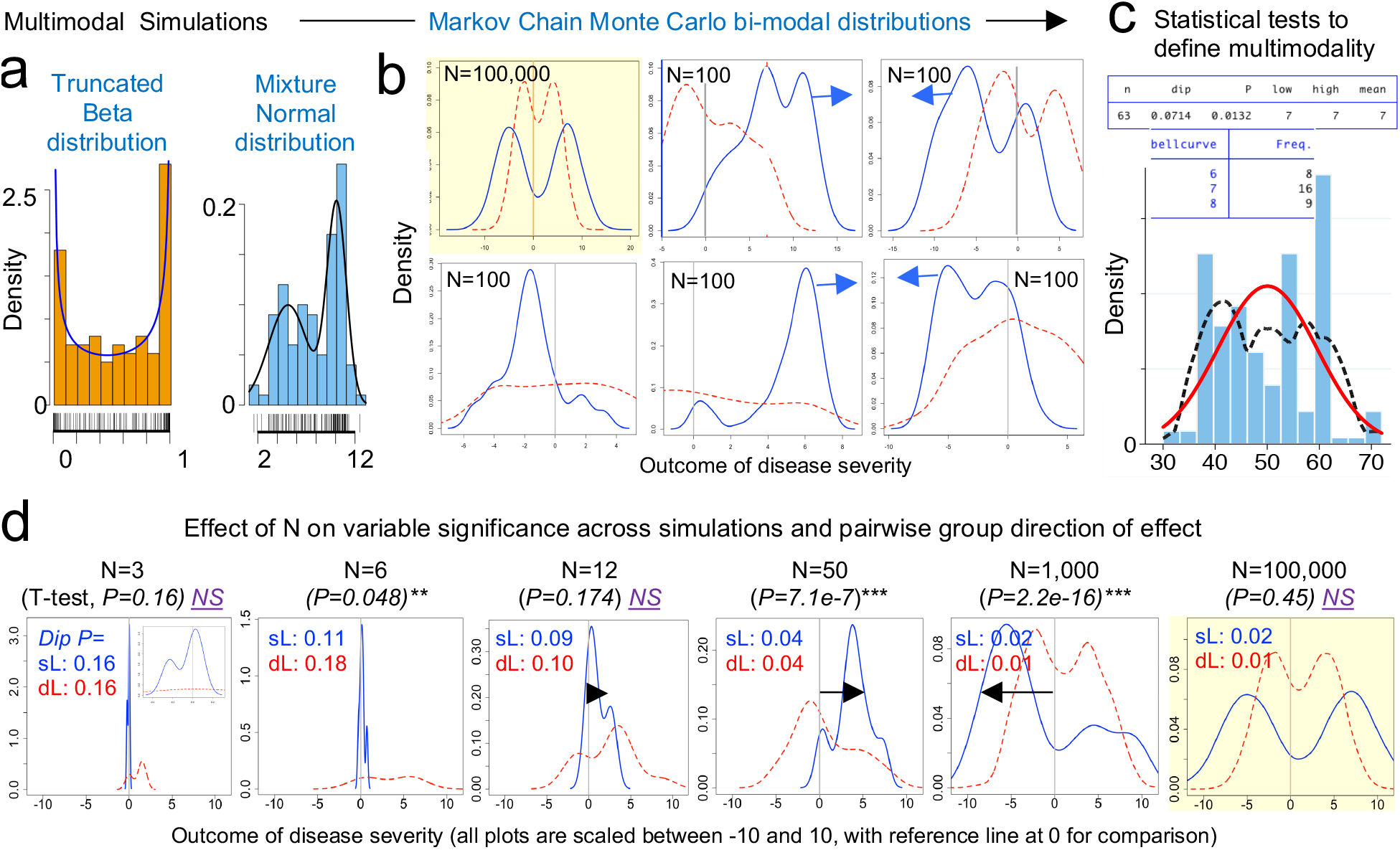
Markov chain Monte Carlo (MCMC) simulations emphasize targeted study of subtypes in the study of multimodal diseases. Random walk Markov chain Metropolis-Hastings’ algorithm to simulate random sampling accounting for the hypothetical dependence of two different disease subtypes. **a)** Mathematical function that allows distribution of number sampling if numbers that follow a bi-modal distribution. Simulation depicts distributions derived from ‘truncated beta’ and the combination of two ‘mixed unimodal’ distribution functions. **b)** Random sampling iterations after Markov chain simulations (N=100) comparing two hypothetical bimodal data distributions (red dotted line vs. blue solid line) for an outcome of disease severity, wherein 100,000 simulations represent approximately the real distribution (grey line; zero). As a stochastic model, the Markov chain algorithm considers biologically relevant sampling dependence. Notice how random sampling for two bimodal distributions can yield non-consistent statistical results variability between iterations, in this case N=100 (−5, SD of 3; 7, SD of 3). **c)** Example of a Hartigan-Hartigan (Hartigans’) unimodality *dip test* and a *modes* test ^61,62^ showing a multimodal data distribution of a hypothetical dataset (black dotted line) compared to a normal univariate density plot (red line). To identify data subtypes (modes), the dip test^61^ computes a p-value to help determine if a data is unimodal or multimodal and does not require *a priori* knowledge of potential multimodality and can be interpreted from its test statistics and P-value (p<0.05 indicates data is not unimodal, p>0.05-1.0 indicating at least one data mode exists in dataset). **d)** Effect of increasing N on t-test significance (direction, arrow), and dip test for MCMC simulations using the Markov chain simulation scripts (panel C) comparing two hypothetical bimodal data distributions (red dotted line vs. blue solid line), controlling for randomness (*set.seed* 101) while increasing N. Plots scaled between −10 and 10 illustrate increased data dispersion as N increases (grey line; zero). See **Supplementary Figure 2** for wider range of N and the examples for dip test and modes analysis using STATA and R commands.

Thus, we used ‘Random walk Markov chain Metropolis-Hastings algorithms’ to simulate random sampling, accounting for the hypothetical dependence between two different disease subtypes. To simulate the statistical comparison of two these two hypothetical bimodal diseases, we ***i)*** ran Markov Chain Monte Carlo (MCMC) simulations (**Figure 5B**), ***ii)*** used the ‘dip test’ to determine if the simulated data were statistically multimodal **Figure 5C**, and ***iii)*** used the Student’s t-test to determine the statistical significance, the mean differences and directions for the simulated distributions, using N=100. The MCMC simulations clearly illustrate how random sampling of two bimodal hypothetical diseases lead to inconsistent patterns of statistical results when compared. Notice that the data dispersion increases as N increases; see summary statistics in **Figure 5D**.

Conclusively, MCMC illustrations emphasize that increasing N in the study of multimodal diseases in a single study should not be assumed to provide results that can be directly extrapolated to the population, but rather, MCMC emphasize that the target study of data subtypes could lead to the identification of mechanisms which could explain why diseases vary within biological systems (*e.g*., humans and mice).

### Personalized ‘data disease subtyping’ must be combined with proper ‘cage-cluster’ statistics

One important caveat to consider across animal studies is that increasing N alone is futile if clustered-data statistics are not used to control for animal cage-density (>1 mouse/cage), which our group showed contributes to ‘artificial heterogeneity’, ‘cyclical microbiome bias’, and false-positive/false-negative conclusions^2,54^. To infer the role of scientific decision on the need for particular statistical methods, we examined the studies reviewed by Walter *et al*.^7^ for ‘animal density’ and ‘statistical’ content (see **Methods**). Of note, only one of the 38 studies (2.6%, 95%CI=0.1-13.8%) used proper statistical methods (mixed-models) to control for cage-clustering^18^. Although on average, studies tested 6.6 patients and 6.4 controls/group (range=1-21), most studies were below the average (65.7%, 25/38, 95%CI=48.6-80.4%), with 14 having <4 donors/group (**Figure 6A**). However, of interest, the number of human donors included in a study was inversely correlated with the number of mice/per donor used in the FMT experiments **Figure 6B**.

**Figure 6.**
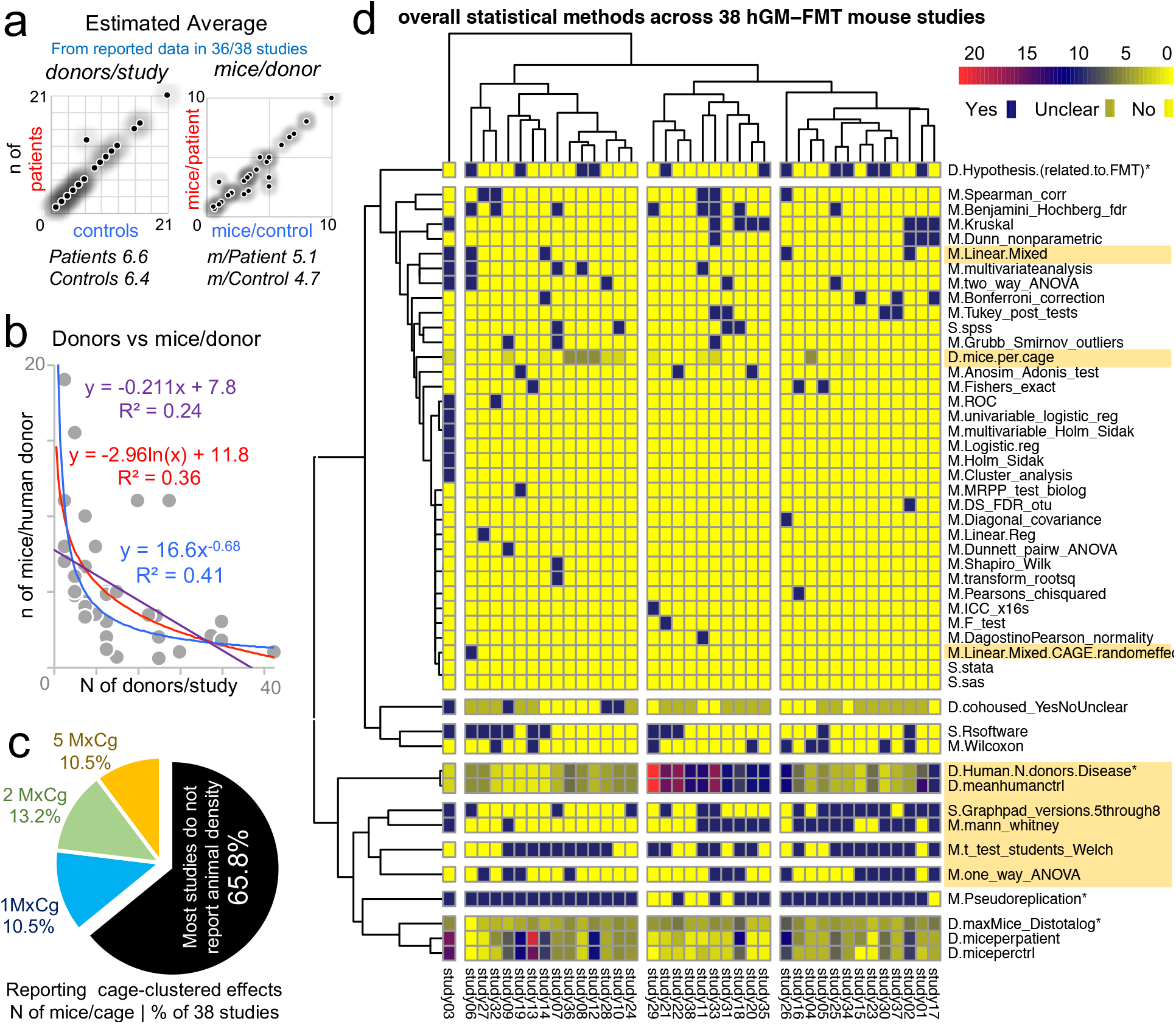
Study design and statistical methods among 38 hGM-FMT studies reveal lack of cage-clustered analysis and dominance of univariate analysis. Analysis of 38 studies as reviewed by Walter *et al*^7^. **a**) Average and correlation for number of donors (left plot; patients with disease vs. healthy control) and average number of mice per human donor in each study (right plot). **b**) Correlation plot with exponential, logarithmic and linear fits shows that scientists tend to use less animals when more donors are tested, creating a ‘trade-off’ between data uncertainty due to variance in human disease with that of variance in animal models for disease of interest. **c**) Pie chart shows distribution of studies reporting mice per cage (MxCg) attributing to cage-clustered effects. (keywords: cage/cluster*, individual/house*, mice per*, density*, mixed/random/fix/methods/stat*, P=). Note that most studies do not report MxCg (*i.e*., animal density). **d**) Heatmap illustrates the overall statistical methods (M), statistical software (S) and study design (D) used by the 38 reviewed studies. Note that only one study (‘study 6’)^18^ used linear mixed methods to control for the random effects of cage clustering and that the majority of studies limited analysis of datasets to univariate-based approaches. *Asterix indicate variables examined by Walter. Notice the dichotomy between software, the GUI interface R statistical software and GraphPad. Highlighted areas shown for reference; notice the cluster around GraphPad and univariate analysis.

Unfortunately, the majority of studies (25/38, 65.8%, 95%CI=48.6-80.4%) did not report animal density, consistent with previous analyses^2^; while 10.5% of the studies (4/38, 95%CI=2.9-24.8%) housed their mice individually, which is advantageous because study designs are free of ICC, eliminating the need for cage-cluster statistics (**Figure 6C**). Our review of the statistical methods used across the 38 studies also revealed that most scientists used GraphPad chiefly for graphics and univariate analysis of mouse phenotype data. This finding suggests an underutilization of the available functions in statistical software, for example, Monte Carlo simulations, to help understand the effect of random sampling on the reproducibility and significance of observed study results, and the likelihood of repeatability by others (Monte Carlo adjusted 95% confidence intervals) (**Figure 6D**).

## DISCUSSION

Despite the inclusion of large numbers of human subjects in microbiome studies, the causal role of the human microbiome in disease remains uncertain. Exemplifying that a large N is not necessarily informative with complex human diseases, a large metanalysis^55^ of raw hGM data from obese and IBD patients showed that human disease phenotypes do not always yield reproducible inter-laboratory predictive biological signatures. Even when hundreds of individuals are studied, especially, if the ‘effect size for the disease of interest’ is narrow (*i.e.,* in obesity; larger in IBD) relative to the variability of the disease. For the human IBD subtypes (*i.e*., ulcerative colitis, and Crohn’s disease), the metanalysis^55^ concluded that only the ileal form of Crohn’s disease showed consistent hGM signatures compared to both healthy control donors and patients with either colonic Crohn’s disease or ulcerative colitis,^56^ but no consistent signatures were observed for obesity. In this context, herein we present observations derived from simulation analysis to highlight that Walter *et al*^7^’s recommendation to scientists to seek further funding to recruit more human donors (increasing N) is an imperfect solution to increase study reproducibly.

Using a simple strategy of assuming random numbers drawn from an observed sample distribution, we have analytically illustrated that increasing N yields aberrant and/or conflicting statistical predictions, which depend on the patterns of disease variability and presence of disease subtypes (data modes). Specifically, our simulations revealed that the number of discernable data subtypes may wax and wane as N increases, and that increasing N does not uniformly enable the identification of statistical differences between groups. Further, subjects randomly selected from a multimodal diseased population may create groups with differences that do not always have the same direction. Especially, ***i)*** if the human disease of interest exhibits variable phenotypes (*e.g*., cancer, obesity, asthma), and ***ii)*** if multivariable cage-clustered data analyses are not used to account for ICC of phenotypes within/between animal cages.

Under the ‘weak law of large numbers’ principle in mathematics (Bernoulli’s theorem^47–49^; see ref for further illustration^57^), as N increases, the distribution of the study/sample means approximates the mean of the actual population, which facilitates the identification of statistically significant (but not biologically meaningful) differences between otherwise overlapping sample datasets. Commonly used statistical methods (*e.g*., t-tests; parametric vs. nonparametric) are designed to quantify differences around the sample centers (mean, median) and range of dispersion (standard errors or deviation) of two groups. However, these methods do not account for the distribution shape (unimodal vs. bi/multimodal) of the compared datasets. With arbitrary increases in N, what is insignificant becomes significant, thus increasing the tendency for the null hypothesis to be rejected despite clinically negligible differences^58,59^.

To guide the selection of sufficient N (cases) or disease data subtype, herein we highlight the use of two simple statistical steps, ***i)*** to first determine if the shape of the dataset is unimodal (*e.g*., dip test), and if not unimodal, then ***ii)*** to use statistical simulations and tests to determine the number of modes/data values of interest. By doing so will facilitate the objective design of personalized/disease subtyping experiments. Although comparisons between group means is important because some diseases are truly different, findings from our own hGM-FMT^46^, and others^16,18^ highlight the relevance of studying disease subtypes and the sources of variability by personalizing the functional analysis of the hGM in mice (*i.e*., that both ‘pathological’ and ‘beneficial’ effects can be seen in hGM-FMT mice independent of donor disease status). For example, in our own work, the functional characterization of ‘beneficial’ or ‘non-beneficial’ disease microbiome subtypes in IBD patients at times of remission could lead to the identification of an ideal patient fecal sample for future autologous transplantation during times of active disease. Therefore, personalized research has the potential to identify different functional microbiome subtypes (on a given outcome, *e.g*., assay or hGM-FMT mice) for one individual.

With respect to determining unimodality, easily implementable tests are available in STATA (*diptest* and *mode*; proprietary and community contributed) and R (Package ‘*multimode*’, community contributed)^60^. The dip test^61^ quantifies departures from unimodality and does not require *a priori* knowledge of potential multimodality and thus information can be easily interpreted from the test statistics and the P-value ^62,63^. Although reports and comparative analysis of statistical performance have been described for various multimodality tests (*e.g*., Dip test, Bimodality test, Silverman’s test and likelihood ratio test^64^, and kernel methods), including simpler alternatives that use benchmarks to determine the influence of data outliers ^52,53,62,65^, it is important to emphasize that every method depends on its intended application and data set (and data shape),^66^ and therefore must be accompanied by the inspection of the data distributions (‘shoulders’, ‘bumps’, and respective ‘valleys’).

In conclusion, by conducting a series of simulations and a review of statistical methods in current hGM-FMT literature, we extensively illustrate the constraints of increasing N as a main solution to identify causal links between the hGM and disease. We also highlight the integral role of multivariable cage-clustered data analyses, as previously described by our group^2^. Herein, we provided a conceptual framework that integrates the dynamics of sample center means and range of dispersion from the compared datasets with kernel and violin plots to identify ‘data disease subtypes’. Biological insights from well-controlled, analyzed and personalized analyses will lead to precise ‘person-specific’ principles of disease, or identification of anti-inflammatory hGM, that could explain clinical/treatment outcomes in patients with certain disease subtypes, and self-correct, guide and promote the personalized investigation of disease subtype mechanisms.

## AUTHOR CONTRIBUTIONS

A.R.P. proposed analytical arguments, conducted statistical analysis and simulations. A.B. and A.R.P conducted the survey, interpreted/analyzed data, and wrote manuscript. F.C. contributed with scientific suggestions and revisions to the manuscript. All authors approved manuscript.

## AVAILABILITY OF DATA AND MATERIALS

All data analyzed for this report are included in the following published articles^7,16,46^ and/or their supplementary information files. The authors will make all statistical codes available upon request.

---

**Supplementary Figure 1.**
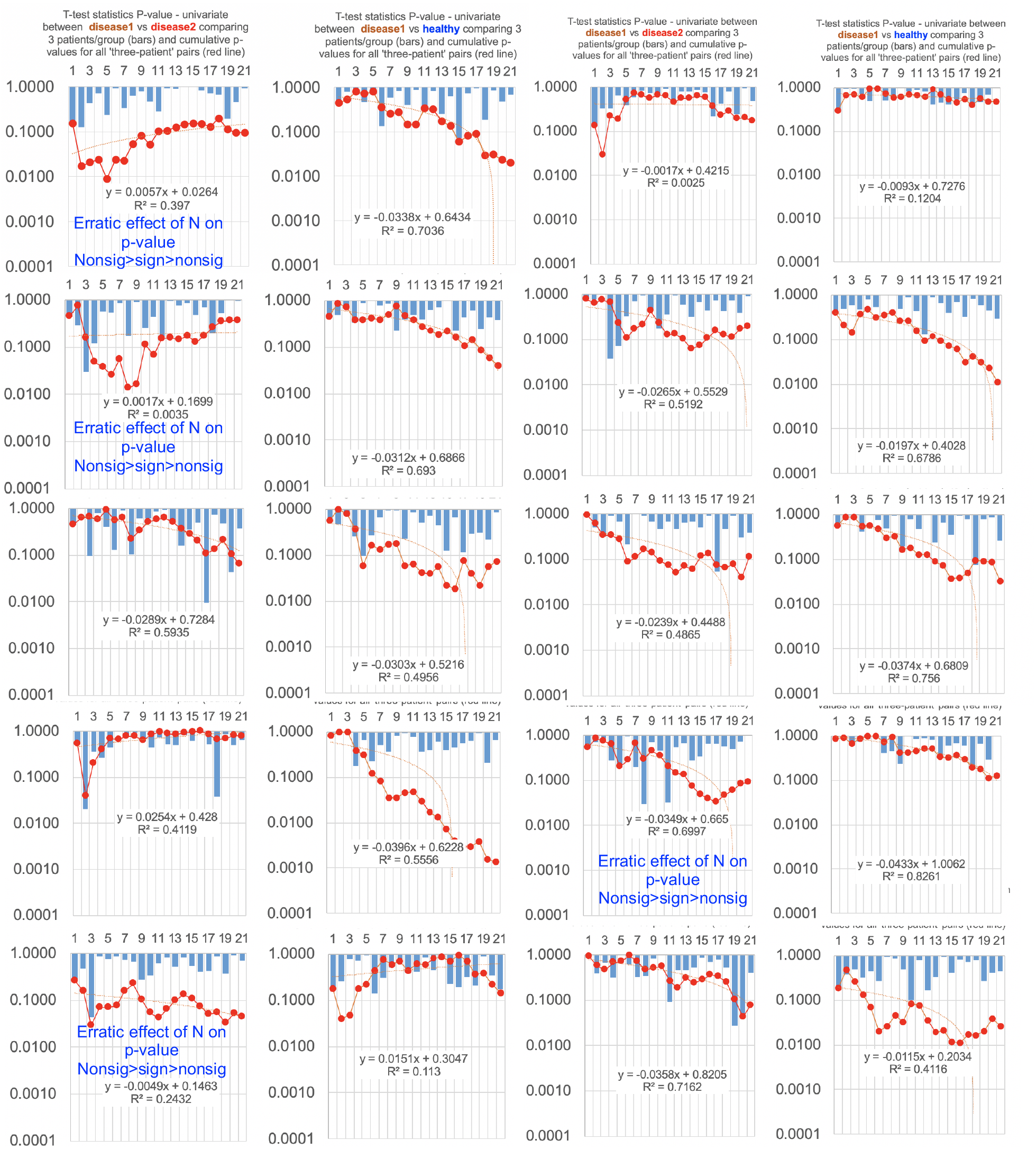
Further examples for data simulations with R^2^ value illustrate linearity as illustrated in Figure 1D. Computed R^2^ value (mean 0.51±0.23, 20 simulations) illustrate the linearity of the correlation between N and statistical significance. Y axis, p-value of the differences using 2-group Student-t test.

**Supplementary Figure 2.**
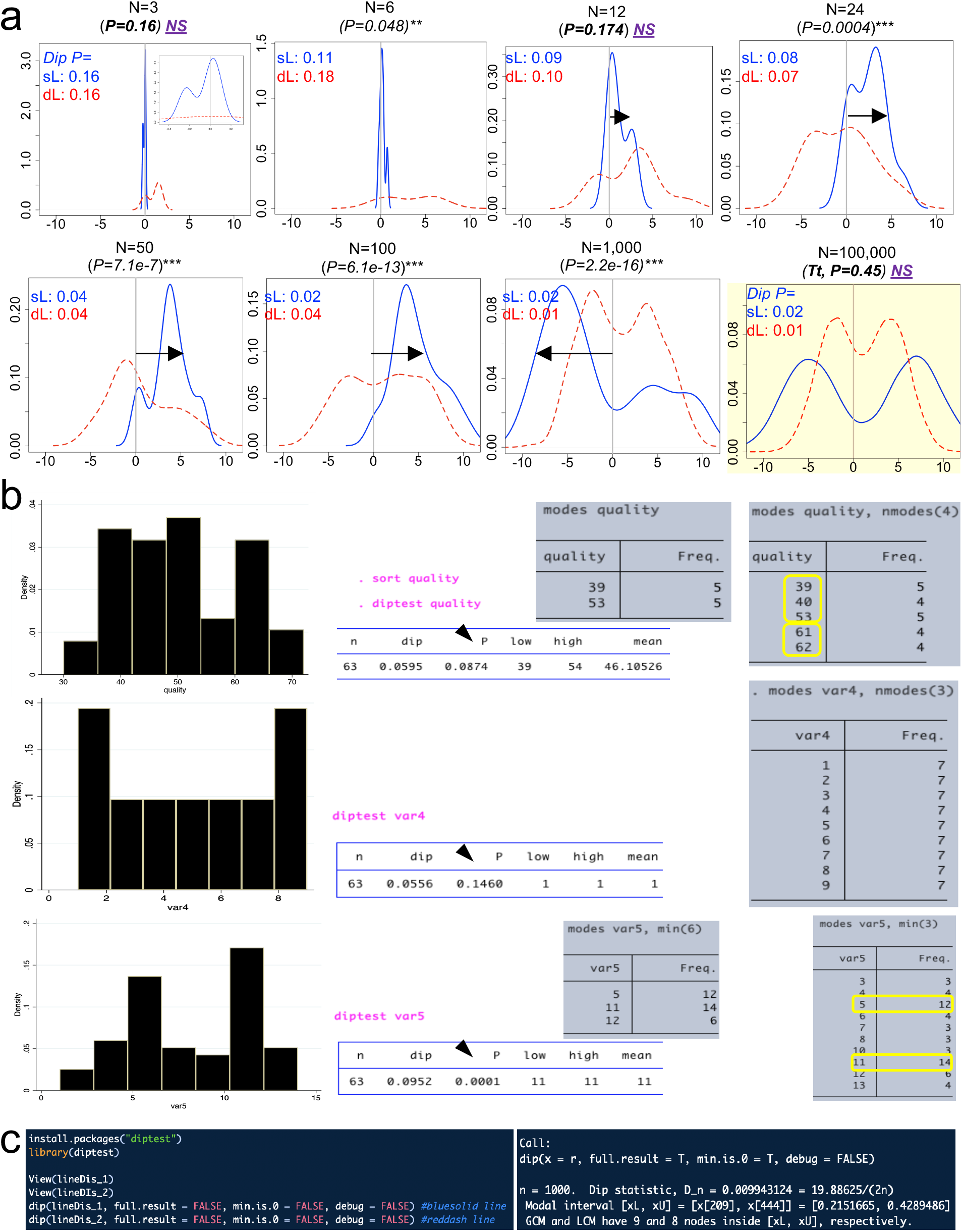
Markov chain Monte Carlo (MCMC) simulations and examples of dip test. Random walk Markov chain Metropolis-Hastings’ algorithm to simulate random sampling accounting for the hypothetical dependence of two different disease subtypes (complement to plots presented in Figure 5).

## METHODS

### Preclinical hGM-FMT (observed) data used for simulations

To facilitate the visualization of how random sampling and disease variability influences study conclusions (significant vs. non-significant p-values) in the context of N, we conducted a series of simulations from preclinical hGM-FMT disease phenotyping data estimates from our own IBD studies (Basson *et al*)^46^ and that of Baxter *et al*^16^ (a study reviewed by Walter *et al*[7]). In brief, by transplanting feces from inflammatory bowel disease (IBD), namely Crohn’s disease (‘Dis1’) and ulcerative colitis (‘Dis2’), and ‘Healthy’ donors (n=3 donors for each ’disease/healthy’ state) into a germ-free spontaneous mouse model of cobblestone/ileal Crohn’s disease (SAMP1/YitFc)^46,67^, Basson *et al*^46^ observed with ~90% engraftment of human microbial taxa after 60 days, that the hGM-FMT effect on mouse IBD-phenotype was independent of the disease state of the donor. Specifically, samples from some IBD patients and some healthy donors did not affect the severity of intestinal inflammation in mice, while the remaining donors exacerbated inflammation. The overlapping presence of both pro-inflammatory and non-inflammatory hGM in the disease phenotype of the mice for IBD and healthy human donors, indicate the presence of data bimodality. Comparably, Baxter *et al*[17] found that differences in the number of tumors resulting in a hGM-FMT mouse model of chemically induced colorectal cancer (CRC) was independent of the cancer status of the human donors (n=3 colorectal cancer, n=3 healthy individuals).

### Simulations from preclinical hGM-FMT data

Simulations were first conducted using random and inverse random normal functions using the mean±SD data from published data^16,46^. In all depicted illustrations, the randomly generated numbers used computer software/automated pseudorandom (seeded and unseeded) methods. Unless described otherwise, the numbers generated were restricted to be confined within biologically meaningful data boundaries based on published data (for example, 0 as minimum for normal histological score or intestinal inflammation, and 80 as arbitrary ~3-fold the maximum possible histological score). For illustration purposes, the x- or y-axes in plots were generically labeled as ‘outcome disease severity’. Simulating a situation where a scientist would recruit a trio of donors (3 donors) per group at a time, and was interested in conducting interim statistical analysis following the addition of every trio of donors to the study, we summarized the pairwise group analysis for the simulated disease comparisons, for various N, and for consecutively added donors as an aggregate ‘cumulative probability of being a significant simulation’ statistic. Comparisons were deemed significant if at least one p-value was <0.05 across simulations. Student’s unpaired t-test and/or One-way ANOVA with Tukey statistical comparisons were also adjusted using Monte Carlo simulations to determine the adjusted p values and the % of simulations to demonstrate that not all disease group comparisons would be reproducible. As a control normal (unimodal) simulation, we created several datasets, including one depicted in illustrations as ‘fake disease X’). Lastly, to illustrate the effect of random sampling from data simulations from multimodal distribution functions, unconstrained-parameter simulations of two mixed yet separate normal distributions, were performed using the Random Walk Metropolis-Hasting algorithm, a form of doing dependent sampling from a proposed posterior distribution, as a well-established method of Markov Chain Monte Carlo (MCMC) simulations, using R, and STATA (v15.1). In the latter, the MCMC sampling of a new individual is dependent on the prior probability of being part of a mode within a bimodal distribution, instead of being completely random from a unimodal distribution, using a log-likelihood correction to prevent negative sigma values and also allow for asymmetrical distributions. This method is beneficial as it asymptotically converges to the true proposal distribution, and so represents a more robust method of data simulation of other potential alternatives of simulating sampling from bimodal distributions (i.e., binomial, and mixed normal distributions).

### Variability of statistical methods in hGM-FMT rodent studies

To determine the sources of statistical methods variability in hGM-FMT rodent studies, we reviewed the content of 38 studies listed in Walter *et al*.^7^ For computation purposes, we searched each article for the following keywords “cage,” “stat*”, “housed,” “multiple,” “multivariable,” “cluster,” “mixed,” “individual*”, “random*” and appropriately extracted details to additional inserted columns of an excel file. Detailed statistical tests and software used, focused on assessing the effect of the hGM in the rodent phenotypes, were extracted to determine if studies used proper cluster statistical analysis, and/or controlled for random effects introduced by caging, when needed; that is, if scientists housed more than one mouse per cage. Integer numbers, including descriptions of animal density, were assigned to the sourced keywords to allow for statistical analysis. If a range was provided for N or animal density (*e.g*., 1-5), estimations were computed using the median value within the range, as well as the minimum and maximum values. Average of estimated center values were used for analysis and graphical summaries

### Statistical Analysis

Descriptive statistics for parametric data were employed if assumptions were fulfilled (*e.g*., 1-way ANOVA). Non-fulfilled assumptions were addressed with nonparametric methods (*e.g*., Kruskal-Wallis). The 95% confidence intervals are reported to account for sample size and for external validity. The test of multimodality was conducted using the dip test (which measures the departure of a sample from unimodality, using as reference the uniform distribution as a worst case) and STATA^61^, with packages available in R^68^. The tabulation of modes from a variable in a dataset was computed using the *modes* and *hsmode* function in STATA.^69,70^. Statistical and simulation analyses were conducted or plotted with Excel, R, Stata, and GraphPad.

